# CpG-adjuvanted stable prefusion SARS-CoV-2 spike protein protected hamsters from SARS-CoV-2 challenge

**DOI:** 10.1101/2021.01.07.425674

**Authors:** Chia-En Lien, Yi-Jiun Lin, Charles Chen, Wei-Cheng Lian, Tsun-Yung Kuo, John D Campbell, Paula Traquina, Meei-Yun Lin, Luke Tzu-Chi Liu, Ya-Shan Chuang, Hui-Ying Ko, Chun-Che Liao, Yen-Hui Chen, Jia-Tsrong Jan, Cheng-Pu Sun, Yin-Shiou Lin, Ping-Yi Wu, Yu-Chiuan Wang, Mi-Hua Tao, Yi-Ling Lin

## Abstract

The COVID-19 pandemic presents an unprecedented challenge to global public health. Rapid development and deployment of safe and effective vaccines are imperative to control the pandemic. In the current study, we applied our adjuvanted stable prefusion SARS-CoV-2 spike (S-2P)-based vaccine, MVC-COV1901, to hamster models to demonstrate immunogenicity and protection from virus challenge. Golden Syrian hamsters immunized intramuscularly with two injections of 1 µg or 5 µg of S-2P adjuvanted with CpG 1018 and aluminum hydroxide (alum) were challenged intranasally with SARS-CoV-2. Prior to virus challenge, the vaccine induced high levels of neutralizing antibodies with 10,000-fold higher IgG level and an average of 50-fold higher pseudovirus neutralizing titers in either dose groups than vehicle or adjuvant control groups. Six days after infection, vaccinated hamsters did not display any weight loss associated with infection and had significantly reduced lung pathology and most importantly, lung viral load levels were reduced to lower than detection limit compared to unvaccinated animals. Vaccination with either 1 μg or 5 μg of adjuvanted S-2P produced comparable immunogenicity and protection from infection. This study builds upon our previous results to support the clinical development of MVC-COV1901 as a safe, highly immunogenic, and protective COVID-19 vaccine.

## Introduction

With over 80 million cases and more than 1.8 million deaths worldwide as of the end of 2020, the COVID-19 pandemic continues to ravage the world one year after its first report in December 2019 [1, 2]. The pandemic also spurred a hitherto unheard of rate of research and vaccine development with 172 vaccines in preclinical development and 61 vaccines in clinical development according to the WHO in December 2020 [3]. The rapid progress of COVID-19 vaccine developed is tracked, for example, by the New York Times’s COVID-19 Vaccine Tracker, which continuously track and update progress of vaccine development and approval [4]. Clearly, the monumental task of controlling this pandemic on a global scale and immunizing a population over 7 billion will require more than a few types of vaccines.

The vast majority of COVID-19 vaccines use the full length or the receptor binding domain of spike (S) protein on the surface of the virus as the antigen, as this binds to human angiotensin converting enzyme 2 (hACE2) for cellular entry and is the major neutralizing antibody inducing antigen [5]. Various modifications including modification of two prolines and inactivation of the furin site have been made to the S protein to lock in its prefusion form to enhance its stability and immunogenicity, and this has been applied to current vaccine development [6-9]. We have previously reported preclinical immunogenicity and safety results of prefusion stabilized S protein, S-2P, adjuvanted with CpG 1018 and aluminum hydroxide (alum) in rodent models [10]. The adjuvanted S-2P (MVC-COV1901) was highly immunogenic and promoted a Th1-biased immune response in mice and no serious adverse effects were observed in toxicology studies in rats [10]. Based on these results, we have carried out the current study in order to investigate the *in vivo* efficacy of MVC-COV1901 in an animal model which is permissive to SARS-CoV-2 and displays symptoms of infection.

Although non-human primates have been used for challenge studies involving SARS-CoV-2 due to similarity of ACE2 receptors and relative closeness to human, the limited availability and high cost are increasingly prohibitive [11]. Small rodent models provide a more economical means of studying the virus; however, mouse ACE2 receptors do not allow permissive infection of SARS-CoV-2 and genetic modification of mice to express human ACE2 (hACE2) or transient transduction using adenovirus-associated virus (AAV) of hACE2 are laborious and costly [12]. Golden Syrian hamsters were found to have the closest homologue of hACE2 and can be infected in lower respiratory tract presenting with symptoms such as weight loss, respiratory distress and lung injury, thus making them an attractive small animal model with which to study SARS-CoV-2 challenge and vaccine development [12-14].

In this study, we present data from a hamster challenge study to test MVC-COV1901 using CpG 1018 and alum adjuvanted S-2P. Potent immunogenicity was induced and hamsters were protected from SARS-CoV-2 infection as demonstrated by the findings that (a) no decreases in body weight were observed in hamsters immunized with both low and high dosage of the vaccine candidate antigen; (b) virus was undetectable in the lungs of immunized hamsters at 3 days after infection by fifty-percent tissue culture infective dose (TCID_50_); and (c) immunized hamsters were protected from lung injury at 6 days after challenge, precluding potential vaccine-associated enhanced respiratory disease (VAERD). These results provide additional evidence for the advancement of our clinical development of MVC-COV1901, of which a phase II trial is current underway (NCT04695652).

## Results

### Hamsters as SARS-CoV-2 virus challenge model for MVC-COV1901

To develop a SARS-CoV-2 virus challenge model in hamsters for MVC-COV1901, an initial study was conducted to determine the optimal dose of virus for the challenge experiments. Unvaccinated hamsters were inoculated with 10^3^, 10^4^, or 10^5^ PFU of SARS-CoV-2 and euthanized on Day 3 or 6 after infection for tissue sampling (Figure S1). Following infection of 10^3^ to 10^5^ PFU of SARS-CoV-2, the hamsters exhibited dose-dependent weight loss. Hamsters infected with 10^3^ PFU gained weight while 10^4^ and 10^5^ PFU-infected hamsters experienced progressively severe weight loss at 6 days post-infection (d.p.i.) (Figure S2). However, there were no significant differences between levels of viral genome RNA (Figure S3a) and viral titer (Figure S3b) measured in 10^3^ to 10^5^ PFU of SARS-CoV-2-infected hamsters at 3 and 6 d.p.i. All dosages of virus resulted in elevated lung pathology (Figure S4), even at 10^3^ PFU where the animals did not experience weight loss (Figure S2). There was also no virus inoculation dose-dependent effect on lung pathology scores and lung viral load (Figures S3, S4). Therefore 10^4^ PFU of virus was used for virus challenge studies as it provides an adequate balance between clinical signs and virus titer for inoculation.

### Administration of S-2P adjuvanted with CpG 1018 and aluminum hydroxide to hamsters induced high levels of neutralizing antibodies

The main study is outlined as in Figure 1: Hamsters were divided into four groups receiving two immunizations at 21 days apart of either vehicle control (PBS only), adjuvant alone, low dose (LD) or high dose (HD) of MVC-COV1901. No differences in body weight changes were observed after vaccination among the four groups (Figure S5). Fourteen days after the second immunization, high level of neutralizing antibody titers were found in both LD and HD groups with ninety-percent inhibition dilution (ID_90_) geometric mean titer (GMT) of 2,226 and 1,783, respectively (Figure 2a). Anti-S IgG antibody levels were high enough that several individual samples reached the upper threshold of detection, with GMTs of LD and HD groups of 1,492,959 and 1,198,315, respectively (Figure 2b). In general, even at low dose, MVC-COV1901 induced potent levels of immunogenicity in hamsters.

**Figure 1.**
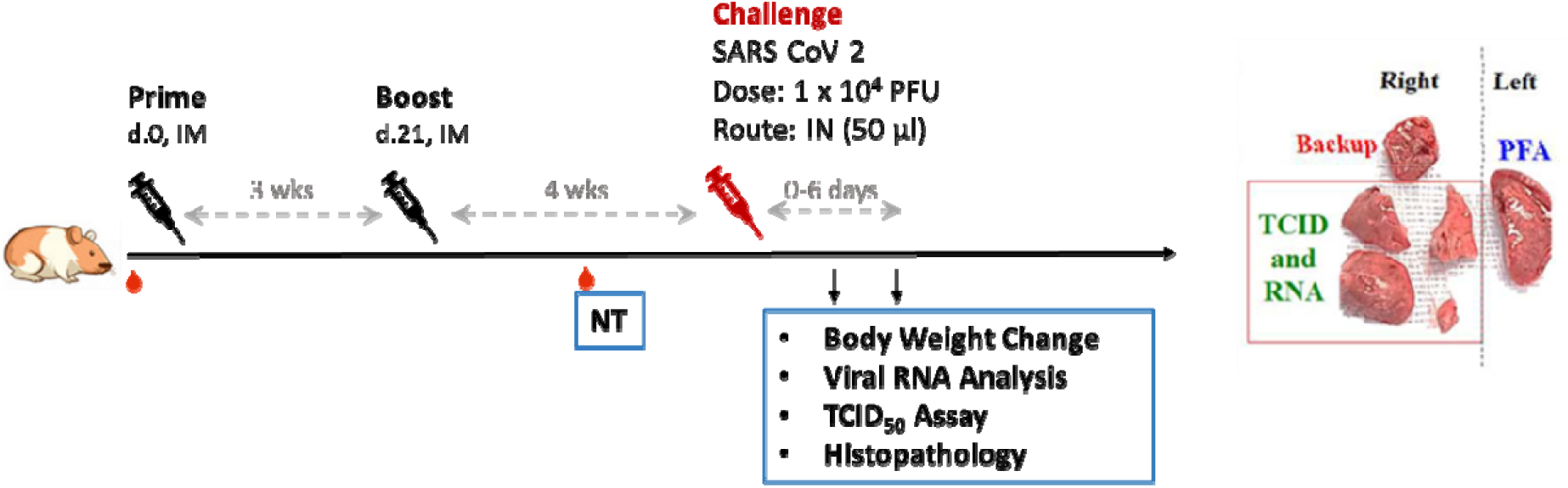
Study design of the hamster challenge study. Hamsters were immunized twice at 3 weeks apart and 2 weeks after the second immunization, serum samples were taken for immunogenicity assays. Four weeks after the second immunization, hamsters were challenged with 10^4^ PFU of SARS-CoV-2. Body weights were tracked for 3 to 6 days after infection and the animals were euthanized on the third or sixth day after infection for necropsy and tissue sampling.

**Figure 2.**
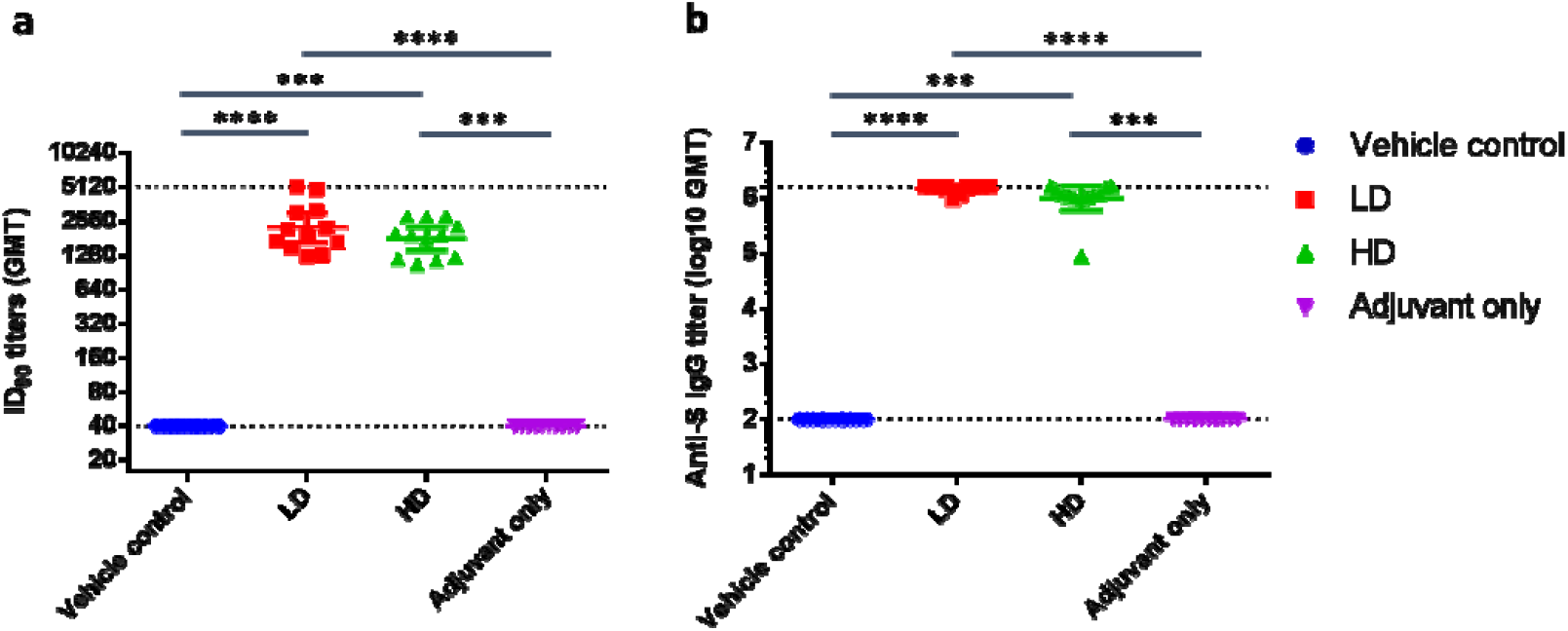
Neutralizing antibody titers with pseudovirus assay in hamsters 2 weeks after second immunization. Hamsters (N=10 per group) were immunized twice at 3 weeks apart with vehicle control (PBS), 1 µg (LD) or 5 µg (HD) of S-2P adjuvanted with 150 µg CpG 1018 and 75 µg aluminum hydroxide, or with adjuvant alone. The antisera were harvested at 2 weeks after the second injection and subjected to **a**. neutralization assay with pseudovirus expressing SARS-CoV-2 spike protein to determine the ID_90_ titers of neutralization antibodies and **b**. total anti-S IgG antibody titers with ELISA. Results are presented as geometric mean with error bars representing 95% confidence interval and statistical significance calculated with Kruskal-Wallis with corrected Dunn’s multiple comparisons test. Dotted lines represent lower and upper limits of detection (40 and 5120 in ID_90_, 100 and 1,638,400 in IgG ELISA).

### Adjuvanted S-2P protected hamsters from clinical signs and viral load after SARS-CoV-2 challenge

Four weeks after the second immunization, hamsters were challenged with 10^4^ PFU of SARS-CoV-2 virus and body weights were tracked up to 3 or 6 days post infection (d.p.i.). Groups of animals were sacrificed on 3 or 6 d.p.i. for viral load and histopathology analyses (Figure 1). LD and HD vaccinated groups did not show weight loss up to 3 or 6 days after virus challenge and instead gained 5 and 3.8 g of mean weight at 6 d.p.i., respectively (Figure 3). The protective effect was most significant at 6 d.p.i. in both vaccinated groups, while vehicle control and adjuvant only groups experience significant weight loss (Figure 3). Lung viral load measured by viral RNA and TCID_50_ assays showed that both viral RNA and viral titer decreased significantly at 3 d.p.i. in vaccinated hamsters and dropped to below the lower limit of detection at 6 d.p.i. (Figure 4). Note that viral load, especially viral titer measured by TCID_50_ dropped noticeably at 6 d.p.i. in control and adjuvant only groups due to hamsters’ natural immune response (Figure 4). Lung sections were analyzed and pathology scoring was tabulated (Figure 5). There were no differences at 3 d.p.i. between control and experimental groups; however, at 6 d.p.i., the vehicle control and adjuvant only groups had significantly increased lung pathology including extensive immune cell infiltration and diffuse alveolar damage, compared to the HD antigen/adjuvant immunized groups (Figure 5, S6). These results showed that MVC-COV1901-induced robust immunity was able to suppress viral load in lungs and prevent weight loss and lung pathology in infected hamsters.

**Figure 3.**
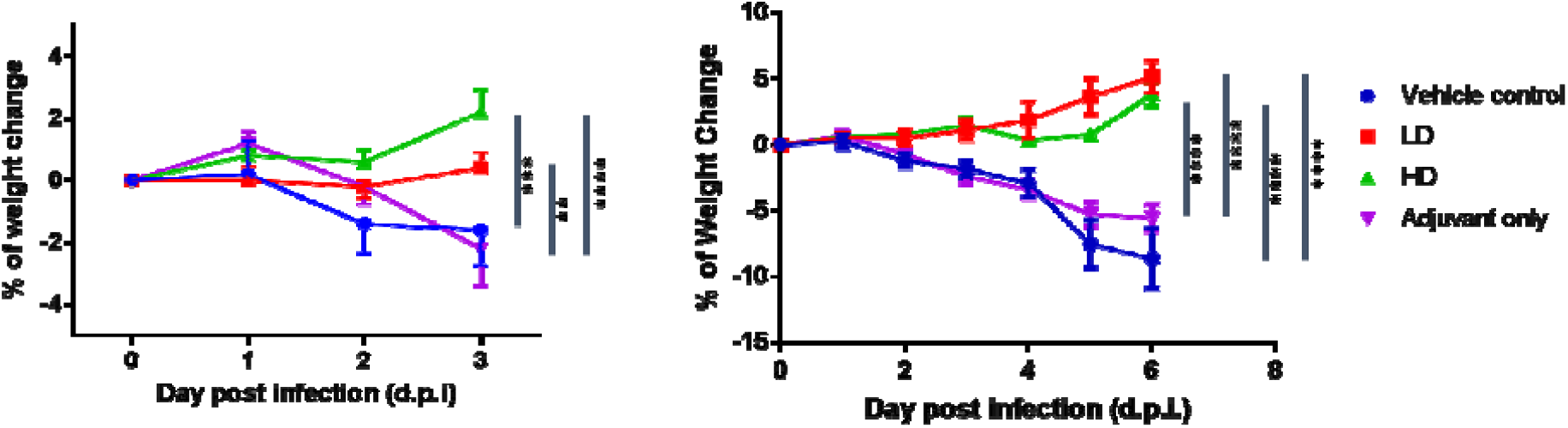
Change in body weight in hamsters after infection with SARS-CoV-2. Hamsters immunized in Figure 2 were challenged with 10^4^ PFU virus. The body weights of individual hamsters were tracked daily up to the time of euthanizing at 3 d.p.i. (n = 5 per group) and 6 d.p.i. (n = 5 per group). Results are presented as mean with error bars representing standard error and statistical significance calculated with Two-way ANOVA with Tukey’s multiple comparison test at 3 d.p.i. (left) or 6 d.p.i. (right).

**Figure 4.**
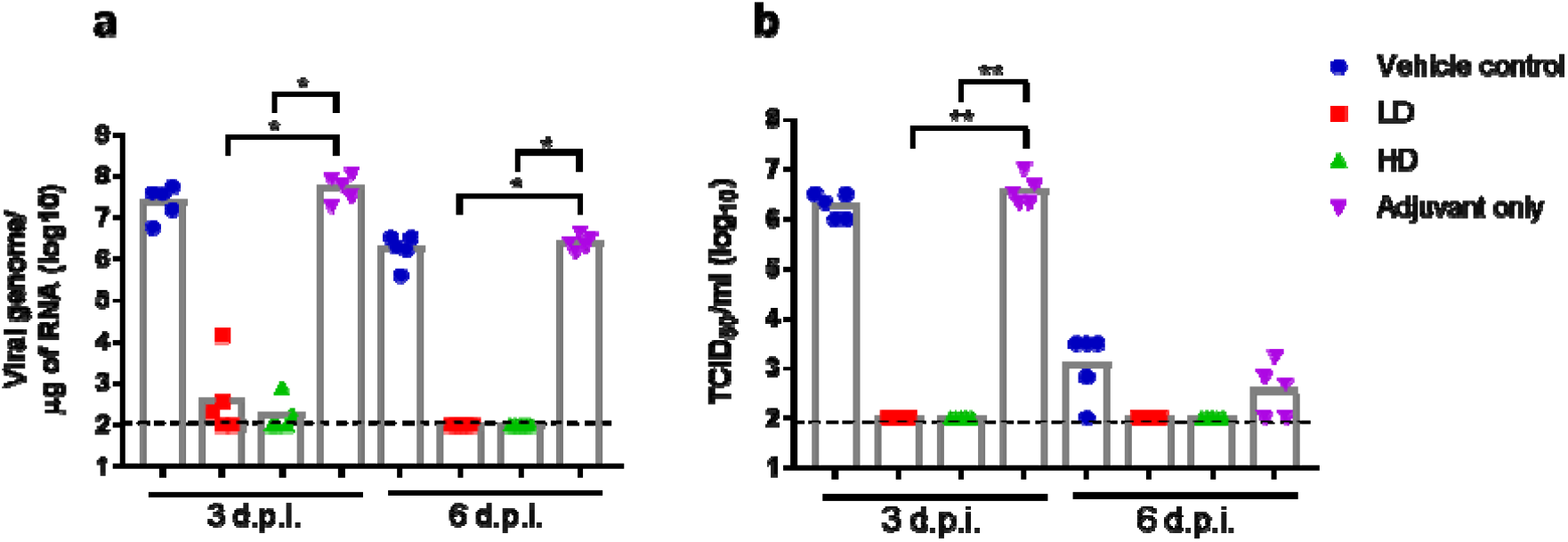
Viral load in hamsters 3 or 6 days post infection with SARS-CoV-2. The hamsters were euthanized at 3 or 6 d.p.i. and lung tissue samples were collected for viral load determination by **a**. quantitative PCR of viral genome RNA, and **b**. TCID_50_ assay for virus titer. Results are presented as geometric mean with error bars representing 95% confidence interval and statistical significance calculated with Kruskal-Wallis with corrected Dunn’s multiple comparisons test. Dotted lines represent lower and limit of detection (100).

**Figure 5.**
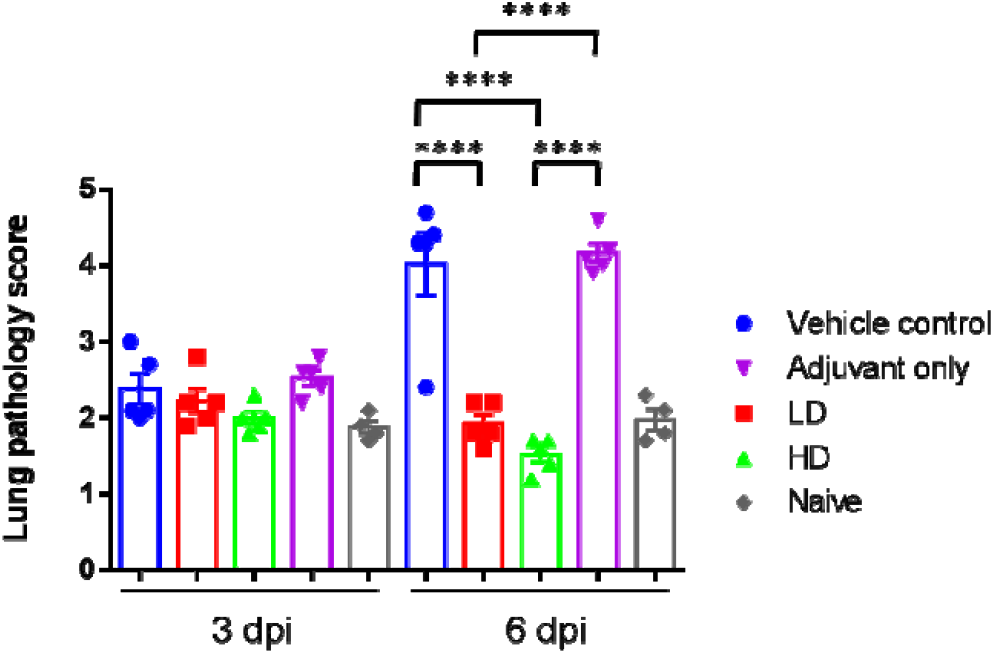
Lung pathology scoring in hamsters 3 or 6 days post infection with SARS-CoV-2. The hamsters were euthanized at 3 or 6 d.p.i. and lung tissue samples were collected for sectioning and staining. The histopathology sections were scored as outlined in the methods and the results tabulated. Results are presented as mean of lung pathology scores with error bars representing standard error and statistical significance calculated with one-way ANOVA with Tukey’s multiple comparisons test.

## Discussion

This report contains the first *in vivo* study that evaluates the preclinical efficacy of MVC-COV1901. A preliminary study helped identify the optimal timing for the observation of change of viral load as measured by viral RNA and infectious virus dose (TCID_50_), which was 3 d.p.i., and 6 d.p.i., respectively. The assays established by Academia Sinica allowed for the observation of a wide window of viral load using both RT-qPCR or TCID_50_. No infectious virus was detected after 3 d.p.i. in hamsters immunized with low dose or high dose of MVC-COV1901, while the low dose arm showed positive for viral RNA at 3 d.p.i.. The discrepancy could be a result of any remaining inoculated virus or virus inactivated by the antibodies. The measurement of sub-genomic RNA (sgRNA) could have helped distinguish the amplifying virus from inactivated virus [15]. All of the hamsters in the MVC-COV1901-immunized groups were protected with significantly reduced lung pathology (generally graded minimal to mild, with a mean score of 1.72 in LD and HD groups), in contrast to diffuse alveolar damage (graded moderate to severe, with a mean score of 4.09 in vehicle and adjuvant control groups) caused by the virus in the lungs of hamsters, in the control groups at 6 d.p.i.. The significance of this study lies not only in the demonstration of *in vivo* efficacy, but also in safety. The viral challenge study allowed for the assessment of risk of disease enhancement with the vaccine candidate. The histopathology scores of the immunized groups have not differed from the non-challenged animals; no evidence of vaccine enhancement was found. Following the consensus made by CEPI and Brighton Collaboration in March 2020, the animal study was run in parallel while Phase I study was cautiously proceeding with careful review of safety data [16]. The vaccines used in this study are from the same batch as the ones used in our Phase I study [17]. The result of this study provides more data that supports progression of the vaccine candidate’s clinical development. There are a few limitations of this study. Firstly, the hamsters were challenged with SARS-CoV-2 at 29 days after the second immunization, a relatively short time that did not allow for the evaluation of the durability of protective antibodies. Secondly, none of the animals died in the pre-test or challenge study within the observation time. Thus, the model is not suitable for the evaluation of severe disease or mortality prevention but, rather, is appropriate for evaluation of the effects of immunization on viral challenge-induced moderate disease. Thirdly, nasal swab was not conducted, thus the study did not evaluate the vaccine’s ability to block viral entry or prevent upper respiratory tract infection. Further studies are needed to evaluate the durability of the protective antibody, the capacity of MVC-COV1901 to prevent severe disease, mortality, or viral entry.

## Methods

### Production of S-2P protein ectodomains from ExpiCHO-S cells

SARS-CoV-2 (Wuhan-Hu-1 strain, GenBank: MN908947) S-2P proteins containing residues 1–1208 with a C-terminal T4 fibritin trimerization domain, an HRV3C cleavage site, an 8×His-tag and a Twin-Strep-tag were produced in ExpiCHO-S cells (ThermoFisher) as described previously [10].

### Pseudovirus-based neutralization assay and IgG ELISA

Lentivirus expressing the Wuhan-Hu-1 strain SARS-CoV-2 spike protein was constructed and the neutralization assay performed as previously described [10]. Briefly, HEK293-hACE2 cells were seeded in 96-well white isoplates and incubated overnight. Sera from vaccinated and unvaccinated hamsters were heat-inactivated and diluted in MEM supplemented with 2% FBS at an initial dilution factor of 20, and then 2-fold serial dilutions were carried out for a total of 8 dilution steps to a final dilution of 1:5120. The diluted sera were mixed with an equal volume of pseudovirus (1,000 TU) and incubated at 37 °C for 1 hour before adding to the plates with cells. Cells were lysed at 72 hours post-infection and relative luciferase units (RLU) was measured. The 50% and 90% inhibition dilution titers (ID_50_ and ID_90_) were calculated referencing uninfected cells as 100% neutralization and cells transduced with only virus as 0% neutralization. Total serum anti-S IgG titers were detected with direct ELISA using custom 96-well plates coated with S-2P antigen.

### Animals and ethical statements

Female golden Syrian hamsters aged 6-9 weeks old on study initiation were obtained from the National Laboratory Animal Center (Taipei, Taiwan). Animal immunizations were conducted in the Testing Facility for Biological Safety, TFBS Bioscience Inc., Taiwan. At 3 weeks after the second immunization, the animals were transferred to Academia Sinica, Taiwan for SARS-CoV-2 challenge. All procedures in this study involving animals were conducted in a manner to avoid or minimize discomfort, distress, or pain to the animals. All animal work in the current study was reviewed and approved by the Institutional Animal Care and Use Committee (IACUC) with animal study protocol approval number TFBS2020-019 and Academia Sinica (approval number: 20-10-1526).

### Immunization and challenge of hamsters

The hamsters were randomized from different litters into four groups (n=10 for each group): hamsters were vaccinated intramuscularly with 2 injections of vehicle control (PBS), 1 or 5 µg of S-2P protein adjuvanted with 150 µg CpG 1018 and 75 µg aluminum hydroxide (alum), or adjuvant alone at 3 weeks apart. The hamsters were bled at 2 weeks after the second immunization via submandibular vein to confirm presence of neutralizing antibodies. Hamsters were challenged at 4 weeks after the second immunization with 1 × 10^4^ PFU of SARS-CoV-2 TCDC#4 (hCoV-19/Taiwan/4/2020, GISAID accession ID: EPI_ISL_411927) intranasally in a volume of 100 µL per hamster. The hamsters were divided into two cohorts to be euthanized on 3 and 6 days after challenge for necropsy and tissue sampling. Body weight and survival rate for each hamster were recorded daily after infection. On days 3 and 6 after challenge, hamsters were euthanized by carbon dioxide. The right lung was collected for viral load determination (RNA titer and TCID_50_ assay). The left lung was fixed in 4% paraformaldehyde for histopathological examination.

### Quantification of viral titer in lung tissue by cell culture infectious assay (TCID_50_)

The middle, inferior, and post-caval lung lobes of hamsters were homogenized in 600 µl of DMEM with 2% FBS and 1% penicillin/streptomycin using a homogenizer. Tissue homogenate was centrifuged at 15,000 rpm for 5 minutes and the supernatant was collected for live virus titration. Briefly, 10-fold serial dilutions of each sample were added onto Vero E6 cell monolayer in quadruplicate and incubated for 4 days. Cells were then fixed with 10% formaldehyde and stained with 0.5% crystal violet for 20 minutes. The plates were washed with tap water and scored for infection. The fifty-percent tissue culture infectious dose (TCID_50_)/mL was calculated by the Reed and Muench method [18].

### Real-time RT-PCR for SARS-CoV-2 RNA quantification

To measure the RNA levels of SARS-CoV-2, specific primers targeting 26,141 to 26,253 region of the envelope (E) gene of SARS-CoV-2 genome were used by TaqMan real-time RT-PCR method described in the previous study [19]. Forward primer E-Sarbeco-F1 (5’-ACAGGTACGTTAATAGTTAATAGCGT-3’) and the reverse primer E-Sarbeco-R2 (5’-ATATTGCAGCAGTACGCACACA-3’), in addition to the probe E-Sarbeco-P1 (5’-FAM-ACACTAGCCATCCTTACTGCGCTTCG-BBQ-3’) were used. A total of 30 μL RNA solution was collected from each lung sample using RNeasy Mini Kit (QIAGEN, Germany) according to the manufacturer’s instructions. Five μL of RNA sample was added into a total 25 μL mixture of the Superscript III one-step RT-PCR system with Platinum Taq Polymerase (Thermo Fisher Scientific, USA). The final reaction mix contained 400 nM forward and reverse primers, 200 nM probe, 1.6 mM of deoxy-ribonucleoside triphosphate (dNTP), 4 mM magnesium sulfate, 50 nM ROX reference dye, and 1 μL of enzyme mixture. Cycling conditions were performed using a one-step PCR protocol: 55°C for 10 min for first-strand cDNA synthesis, followed by 3 min at 94°C and 45 amplification cycles at 94°C for 15 sec and 58°C for 30 sec. Data was collected and calculated by Applied Biosystems 7500 Real-Time PCR System (Thermo Fisher Scientific, USA). A synthetic 113-bp oligonucleotide fragment was used as a qPCR standard to estimate copy numbers of the viral genome. The oligonucleotides were synthesized by Genomics BioSci and Tech Co. Ltd. (Taipei, Taiwan).

### Histopathology

The left lung of hamsters was isolated and fixed in 4% paraformaldehyde. After fixation with 4% paraformaldehyde for one week, the lung was trimmed, processed, embedded, sectioned, and stained with Hematoxylin and Eosin (H&E), followed by microscopic examination. The lung section was evaluated with a lung histopathological scoring system described below [20, 21]:

Lung section is divided into 9 areas and numbered as in the example below:

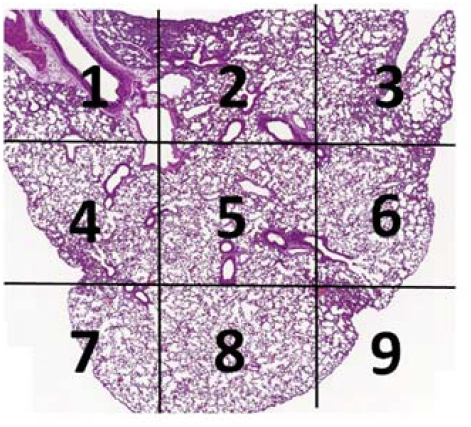

Lung tissue of every area is scored using the following scoring system in the Table 1.

**Table 1.** Lung histopathology scoring system

| Score | Observations |
| --- | --- |
| **0** | Normal, no significant finding |
| **1** | Minor inflammation with slight thickening of alveolar septa and sparse monocyte infiltration |
| **2** | Apparent inflammation, alveolus septa thickening with more interstitial mononuclear inflammatory infiltration |
| **3** | DAD*, with alveolus septa thickening, and increased infiltration of inflammatory cells |
| **4** | DAD, with extensive exudation and septa thickening, shrinking of alveoli, restricted fusion of the thick septa, obvious septa hemorrhage and more cell infiltration in alveolar cavities |
| **5** | DAD, with massive cell filtration in alveolar cavities and alveoli shrinking, sheets of septa fusion, and hyaline membranes lining the alveolar walls |

\*DAD = Diffuse alveolar damage

The average of scores of these 9 areas is used to represent the score of the animal.

### Statistical analysis

The analysis package in Prism 6.01 (GraphPad) was used for statistical analysis. One-way and two-way ANOVA with Tukey’s multiple comparison test and Kruskal-Wallis with corrected Dunn’s multiple comparisons test were used to calculate significance as noted in respective figure descriptions. * = p < 0.05, ** = p < 0.01, *** = p < 0.001, **** = p < 0.0001

## Acknowledgements

We are grateful for the participation of Dr. Michael D. Malison and Dr. Robert L. Coffman for manuscript review and constructive comments, and Wendy Li for organizing the lung histopathology figures. We also thank team members at TFBS Bioscience Incorporation for hamster housing and immunization process. We thank the Biosafety Level 3 Facility, Academia Sinica, Taiwan, for providing environment to handling and performing wild-type virus assay and Dr. Yu-Chi Chou at the RNAi Core Facility, Academia Sinica for the pseudovirus neutralization assay.

## Author Contributions

T.-Y. K. produced the S-2P antigens used in the study. C.-E. L., Y.-J. L., J. D. C., P. T., M.-Y. L., M.-H. T., and Y.-L. L. designed the study and experiments. C.-E. L., Y.-J. L., H.-Y. K., C.-C. L., Y.-H. C., J.-T. J., C-.P. S., Y.-S. L., P.-Y. W., and Y.-C. W. performed and analyzed the experiments. M.-Y. L., L. T.-C. L., and Y.-S. C. drafted the manuscript. All authors reviewed and approved of the final version of the manuscript.

## Data Availability

The datasets generated during and/or analyzed during the current study are available from the corresponding author on reasonable request.

