## Supplementary data for "CpG-adjuvanted stable prefusion SARS-CoV-2 spike protein protected hamsters from SARS-CoV-2 challenge" (from another version of this preprint: version 1, 2021-01-07)

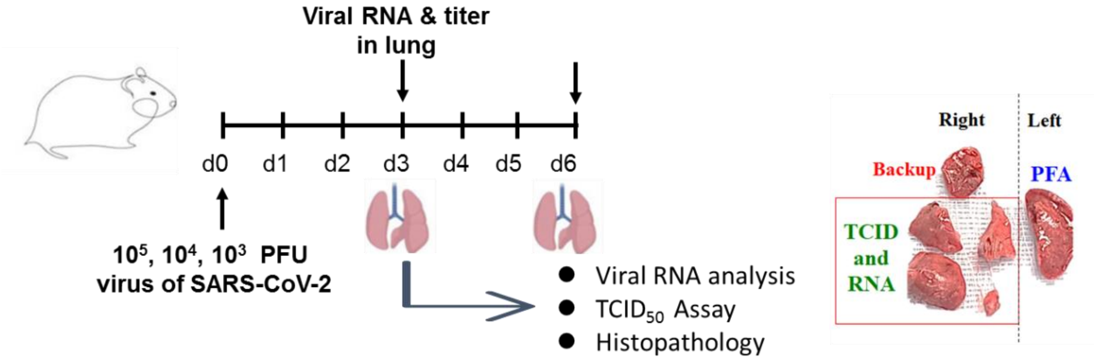


**Figure S1. Study design for hamster challenge model development study.**

Hamsters were challenged with $10^3$, $10^4$, or $10^5$ PFU of SARS-CoV-2. Body weights were measured each day for up to 6 days after infection and hamsters were euthanized on the third and sixth day after infection for necropsy and tissue sampling.

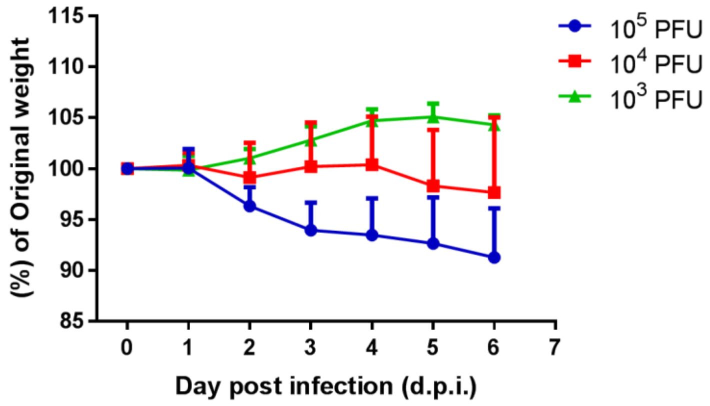


**Figure S2. Body weight change in unvaccinated hamsters after the SARS-CoV-2 infection.**

Body weights of hamsters were monitored each day after virus challenge for 6 days until euthanization. Results are presented as mean with error bars representing standard error.

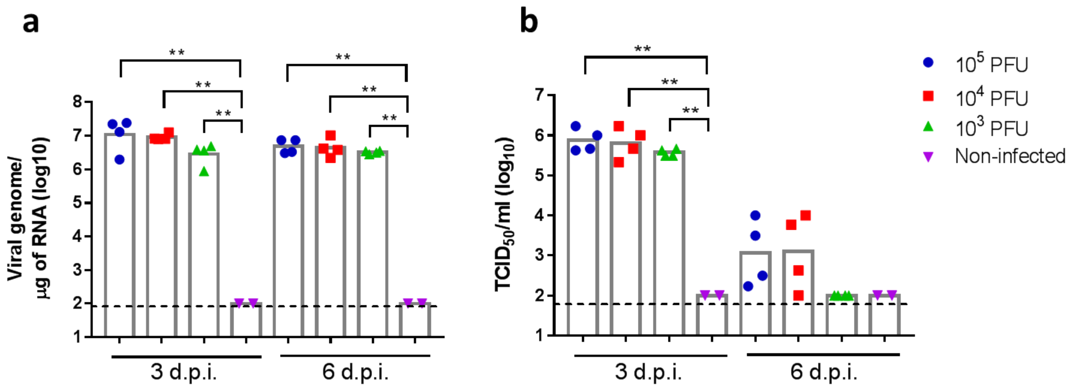


**Figure S3. Viral load in unvaccinated hamsters after the SARS-CoV-2 infection.**

The hamsters were euthanized at 3 or 6 d.p.i. and lung tissue samples were collected for viral load determination by **a**. quantitative PCR of viral RNA, and **b**. $TCID_{50}$ assay for virus titer. Results are presented as geometric mean with error bars representing 95% confidence interval and statistical significance calculated with Kruskal-Wallis with corrected Dunn's multiple comparisons test. Dotted lines represent lower and limit of detection (100).

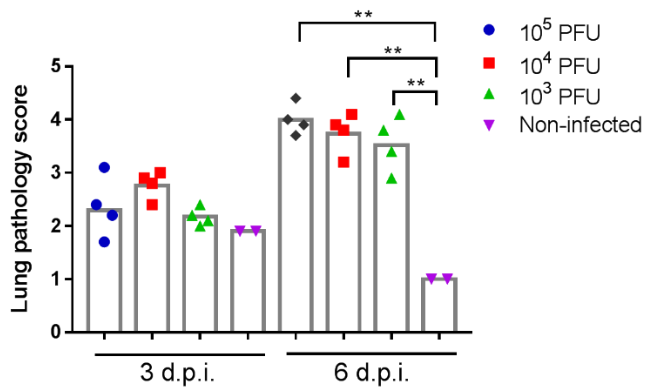


**Figure S4. Lung pathology scores in unvaccinated hamsters after the SARS-CoV-2 infection.**

The hamsters were euthanized at 3 or 6 d.p.i. and lung tissue samples were collected for sectioning and staining. The histopathology sections were scored as outlined in the methods and the results tabulated. Results are presented as mean of lung pathology scores with error bars representing standard error and statistical significance calculated with Kruskal-Wallis with corrected Dunn's multiple comparisons test between groups.

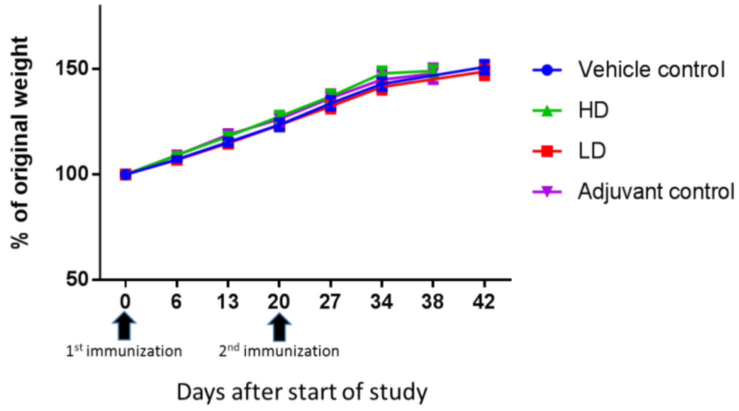


**Figure S5. Body weight change in hamsters after start of the study.**

Body weights of hamsters were monitored on days 0, 6, 13, 20, 27, 34, 38 (LD and adjuvant only), and 42 (vehicle control and HD) after the first immunization. Results are presented as mean with error bars representing standard error.

**Figure S6. Histopathology sections in individual non-infected and infected hamsters in the MVC-COV1901 immunized and control groups.**

Hamsters were divided into a non-infected group (n = 8) and 4 infected groups (n = 5 per group) which had received two immunizations at 21 days apart of either vehicle control, adjuvant alone, low dose and high dose of MVC-COV1901. The hamsters were euthanized at 3 or 6 days after SARS-CoV-2 infection and lung tissue samples were collected for sectioning and H&E staining. The representative images of histopathology in lungs from infected and non-infected control group are displayed.

| | |
|---|---|
| **1/ TA-101, Vehicle control, sacrificed D3, #521**<br>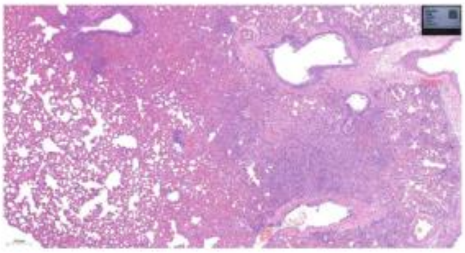 | **1/ TA-101, Vehicle control, sacrificed D3, #522**<br>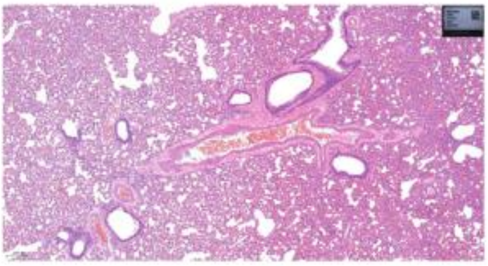 |
| **2/ TA-102, Low dose, sacrificed D3, #526**<br>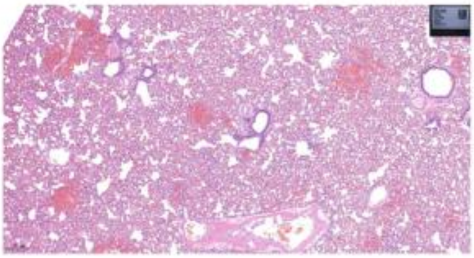 | **2/ TA-102, Low dose, sacrificed D3, #527**<br>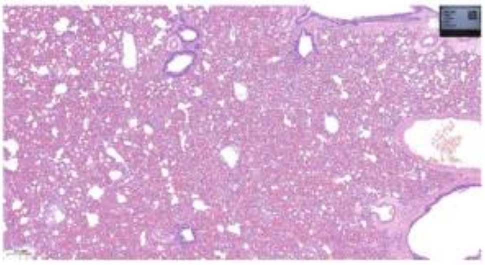 |
| **9/ TA-103, High dose, sacrificed D3, #549**<br>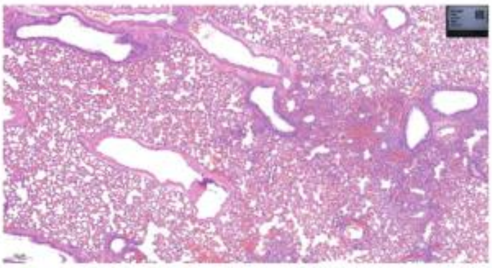 | **9/ TA-103, High dose, sacrificed D3, #550**<br>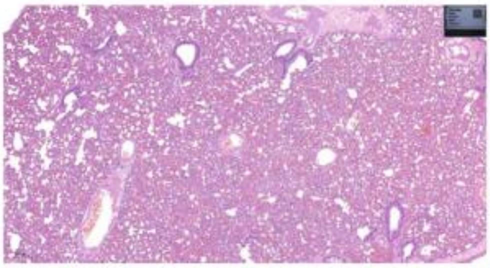 |
| **10/ TA-104, Adjuvant control, sacrificed D3, #554**<br>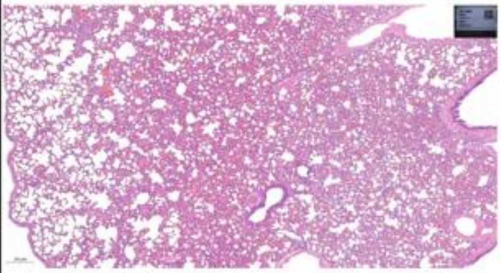 | **10/ TA-104, Adjuvant control, sacrificed D3, #555**<br>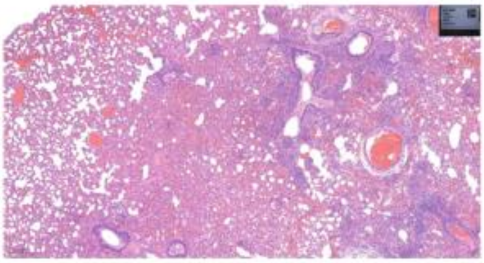 |

**3/ TA-101, Vehicle control, sacrificed D6, #531**

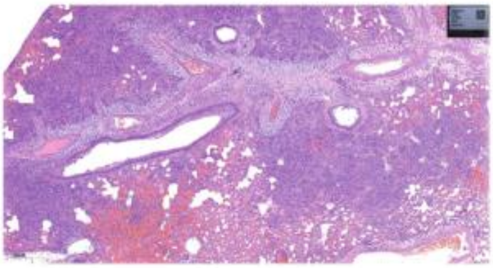

**3/ TA-101, Vehicle control, sacrificed D6, #532**

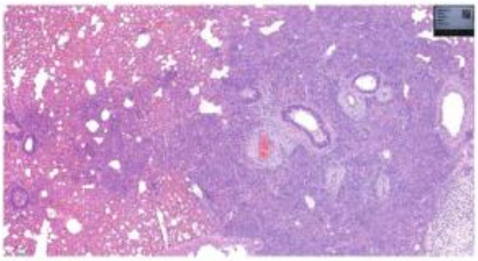

**4/ TA-102, Low dose, sacrificed D6, #536**

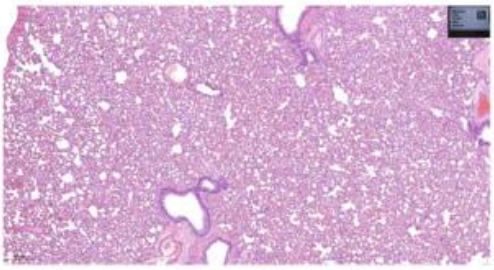

**4/ TA-102, Low dose, sacrificed D6, #537**

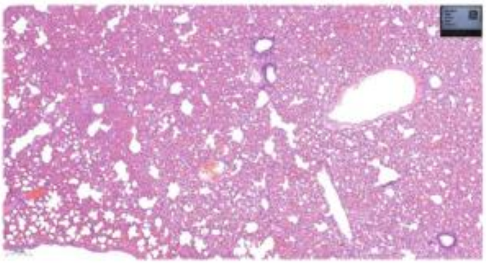

**11/ TA-103, High dose, sacrificed D6, #559**

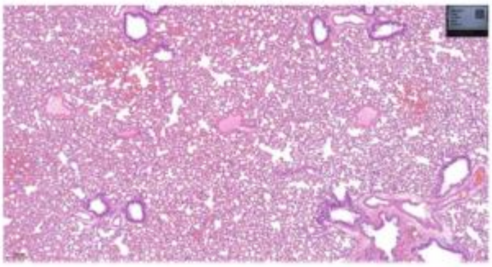

**11/ TA-103, High dose, sacrificed D6, #560**

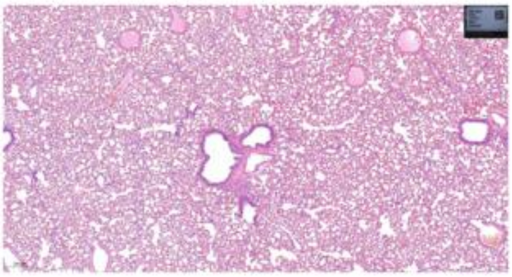

**12/ TA-104, Adjuvant control, sacrificed D6, #564**

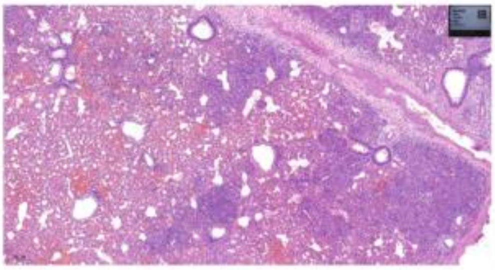

**12/ TA-104, Adjuvant control, sacrificed D6, #565**

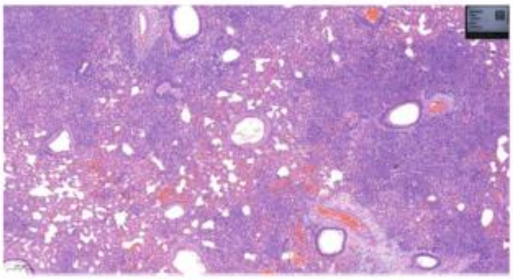

**5/ TA-101, non-infected**

**sacrificed D3, #541**

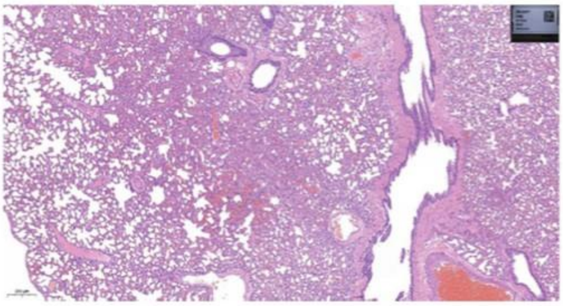

**5/ TA-101, non-infected**

**sacrificed D3, #542**

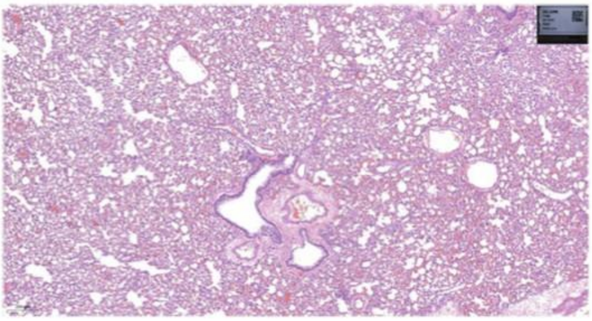

**8/ TA-103, non-infected**

**sacrificed D6, #547**

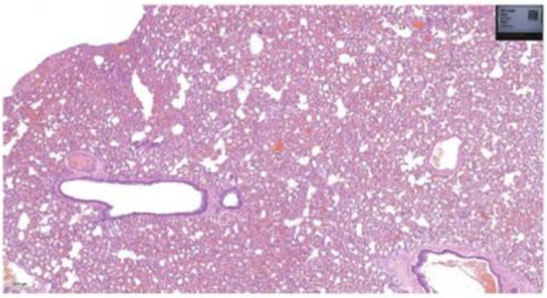

**8/ TA-103, non-infected**

**sacrificed D6, #548**

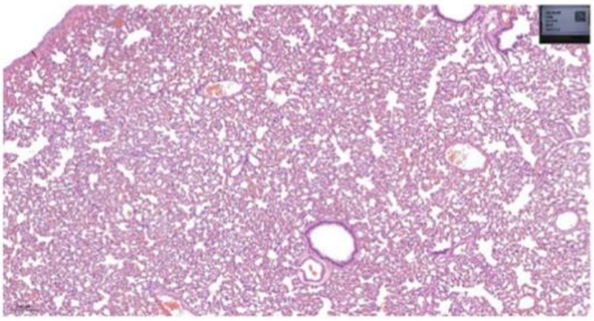
